# Interchromosomal interaction of homologous Stat92E alleles regulates transcriptional switch during stem-cell differentiation

**DOI:** 10.1101/2021.11.08.467622

**Authors:** Matthew Antel, Madona Masoud, Romir Raj, Ziwei Pan, Sheng Li, Barbara G. Mellone, Mayu Inaba

## Abstract

The strength of pairing of homologous chromosomes differs in a locus-specific manner and is correlated to gene expression states. However, the functional impact of homolog pairing on local transcriptional activity is still unclear. *Drosophila* male germline stem cells (GSCs) constantly divide asymmetrically to produce one GSC and one differentiating gonialblast (GB). The GB then enters the differentiation program in which stem cell specific genes are quickly downregulated. Here we demonstrate that a change in local pairing state of the Stat92E locus is required for the downregulation of the Stat92E gene during differentiation. Using OligoPaint fluorescent in situ hybridization (FISH), we show that the interaction between homologous regions of Stat92E is always tight in GSCs and immediately loosened in GBs. When one of the Stat92E locus was absent or relocated to another chromosome, Stat92E did not pair and failed to downregulate, suggesting that the pairing is required for switching of transcriptional activity. The defect in downregulation of Stat92E was also observed upon knockdown of global pairing or anti-pairing factors. Moreover, the Stat92E enhancer element, but not cis-transcription, is required for the change in pairing state, indicating that it is not a consequence of transcriptional changes. GSCs are known to inherit pre-existing histones H3 and H4, while newly synthesized histones are distributed in GBs. When this histone inheritance was compromised, the change in Stat92E pairing did not occur, suggesting that it is an intrinsically programmed process during asymmetric stem cell division. We propose that the change of local pairing state may be a common process to reprogram gene activity during cell-differentiation.

## Introduction

Distant DNA regions interact not only *in cis* but also *in trans* to modify each other’s gene expression states [1-3]. One fascinating facet of inter-chromosomal interaction occurs between homologous chromosomes, the phenomenon called homologous chromosome pairing. Although homologous chromosome pairing is most prominently studied in the context of meiosis, somatic cells of *Dipteran* including *Drosophila* also pair their homologs in somatic cells, across the entire genome and throughout development [4-8]. While the prevalence of complete pairing of the homologs outside of the germline in other organisms is still unclear, somatic pairing of specific chromosome regions does occur in a tightly regulated manner in many other systems including mammals (reviewed in [8]).

Haplotype-resolved Hi-C and/or fluorescent in situ hybridization (FISH) analyses have started to unveil that homologous chromosome pairing has more global impact on 3D genome architecture and gene expression status than previously thought. Local allelic pairing of a particular gene locus differs in a tissue-specific manner [9] and correlates with local chromatin status [10, 11]. This suggests that somatic homolog pairing may be under the control of a developmental program or extracellular signaling. However, the causal relationships between pairing and gene transcription is still uncertain.

How can the interaction of homologous chromosomes influence their transcriptional status? In flies, the consequence of somatic homolog pairing is represented by the phenomenon called transvection, whereby different mutant alleles of a gene-regulatory element can rescue each other’s expression [12-17]. Transvection has been described for a number of *Drosophila* genes and can either promote or silence transcription (reviewed by [14]). Homolog pairing occurs between “buttons” characterized by topology associated domains (TADs) spanning about 100Kb∼ of chromatin domains which can be visualized by Hi-C [9], whereas transvection requires smaller DNA elements such as polycomb responsive elements (PREs) and insulator domains [9] [14]. These requirements account for a recent report showing pairing is necessary but not sufficient for transvection to occur [9]. Even though the phenomenon of transvection was first reported over 60 years ago, its mechanism is still not fully understood. Moreover, because the majority of transvection studies are transgene-based, whether endogenous wild-type genes require trans-homolog interaction to modify their expression level is unclear.

*Drosophila* male germline stem cells (GSCs) constantly divide asymmetrically to produce one GSC and one gonialblast (GB) (Fig. 1A). The GB initiates the differentiation program to enter 4 rounds of syncytial divisions of spermatogonia (SG) then become spermatocytes (SCs). SCs enter 2 rounds of meiotic divisions and ultimately differentiate into functional sperm. Upon exit from GSC state, key stem cell specific genes must be downregulated and genes required for differentiation need to be turned on. Studies have investigated the extrinsic signals and intrinsic factors required for proper cell fate transition during the asymmetric division of GSCs (reviewed in [18]). Particularly, a niche ligand, Unpaired (Upd), has been believed to be a major factor to dictate stem cell fate as ectopic expression of Upd induces overproliferation of GSC-like cells outside of the niche [19]. On the other hand, a number of studies have identified the need of intrinsic factors (reviewed by [20]). For example, newly synthesized histone H3 is inherited by the GB during asymmetric division, while the old histone H3 remains in the GSC [21]. Perturbation of this asymmetric histone H3 inheritance results in differentiation defects [22], demonstrating the indispensability of cell-intrinsic mechanisms. However, it is unclear how such mechanisms collaborate with each other to successfully produce cells with distinct cell fates. Moreover, “ON” and “OFF” timing of key regulatory genes during the fate transition have yet to be elucidated.

**Fig. 1.**
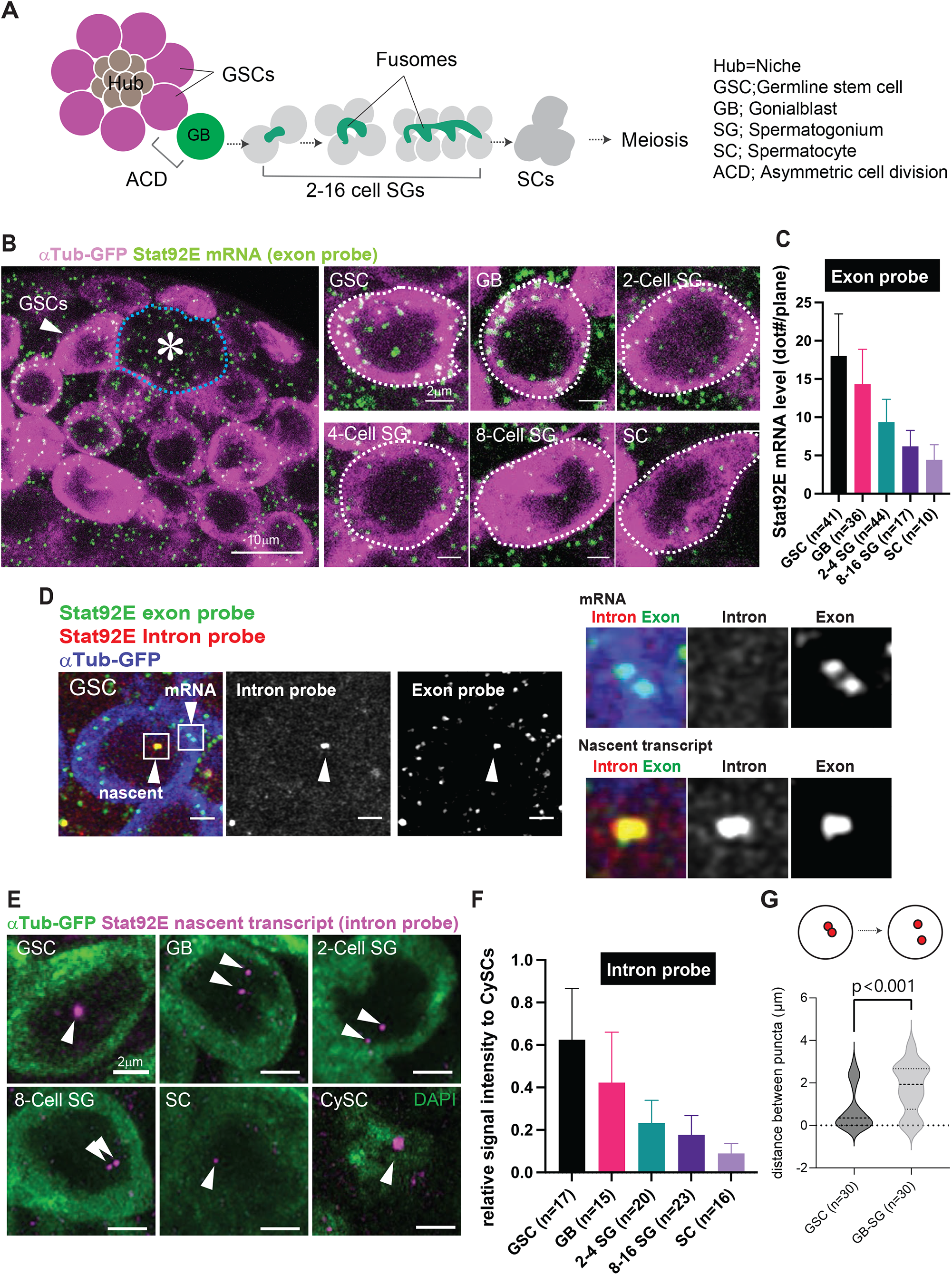
STAT92E transcription level rapidly decreases during differentiation. **A**) Schematic of the *Drosophila* testicular niche and the path of germline differentiation. **B**) Representative images of single molecule RNA FlSH (smFISH) using a Stat92E exon probe (green, see Method for details). Each dot represents a single Stat92E mRNA molecule. αTubulin-GFP (αTub-GFP, magenta) expression under the control of germline driver nosGal4 was used to identify germ cells. The asterisk denotes the hub and entire hub area is encircled by a blue dotted? line. Approximately middle plane of germ cells at the indicated stages are shown in the panels on the right. White dotted outlines encircle the entire cell area from where number of mRNA molecules (dots) were manually counted. **C**) Quantification of Stat92E mRNA in cells at the indicated stages. Y axis values are the average number of smFISH dots present in a middle plane of the cell (dot#/plane) as shown in **B. D**) Representative FISH image showing nascent Stat92E transcripts in GSC. Right panels show magnified regions from insets in the left column. Nascent transcripts are detected by both an intron probe (red) and an exon probe (green) in the nucleus. mRNA signals in the cytoplasm are positive for the exon probe but not the intron probe. Germ cells are visualized by nos>αTubulin-GFP (blue). **E**) Representative RNA FISH images visualizing the nascent transcript of Stat92E (magenta, pointed by white arrowheads) in the cells at the indicated stages. Germ cells are visualized by nos>αTubulin-GFP (green). **F**) Quantification of the amount of Stat92E nascent transcript throughout differentiation. Y axis indicates the average fluorescence intensity of nascent transcript signal in germ cells divided by CySCs’ (see Method for more details). **G**) Violin plots show the distances between Stat92E nascent transcript puncta in GSCs vs. GB-SGs. Cells with more than three puncta were omitted from scoring (see description about Fig 1H). KDE and quantile lines are shown. The width of each curve corresponds with the approximate frequency of data points. Scale bars represent 2μm unless denoted otherwise.

## Results

### STAT92E transcription level rapidly decreases during differentiation

Stat92E is a direct downstream molecule of the niche Upd signal and is known to be required for GSC establishment and maintenance [19, 23-26]. Stat92E protein is specifically expressed in the GSC population and decreases its level immediately in differentiating daughter gonialblasts (GBs) ([27], Fig. S1). It is thought that the GSC-specific Stat92E expression pattern might be regulated by a niche-derived Unpaired (Upd) signal based on a report of mammalian homolog STAT3 [28] in which activated STAT3 protein binds to its own promoter to induce expression, which forms a positive-feedback loop [29]. To test whether the steep gradient of Stat92E expression pattern is regulated at the transcriptional level as hypothesized, we performed single molecule fluorescent *in situ* hybridization (FISH) (smFISH) [30-35]. smFISH visualizes Stat92E mRNA as uniform size/intensity of “dots” in the cytoplasm of germline cells and neighboring somatic cyst stem cells (CySCs) (Fig. 1B). By counting the number of dots in a single confocal plane, we found that Stat92E downregulation occurs gradually as differentiation proceeds from the stage of GSC to SC (Fig. 1B-C).

Although the smFISH results provided precise quantification of Stat92E mRNA, the mRNA can be diluted out upon cell division, leaving open the possibility that the change of transcription occurred even earlier in differentiation. To determine when Stat92E transcription turns off, we designed probes targeting an intron of the Stat92E gene to monitor the level of nascent transcript (Fig. 1D), which represents active transcription occurring on the template DNA region [32, 36, 37] (seen as puncta double positive for exon and intron probe signal, while mRNA shows only exon probe signal, Fig.1E). Quantification of intron signal intensity relative to that in CySCs, which show consistently high signal in a single focus, indicates that the transcription of Stat92E also decreases gradually as differentiation proceeds, with a similar timing to the decrease in mRNA level (Fig. 1F). These results indicate that Stat92E is actively downregulated at its transcriptional level during differentiation from GSC to SC.

Our data indicated that the Stat92E gene is an ideal model gene to study how the transcription of a stem cell gene is downregulated during differentiation. It should be noted that the timing of downregulation of Stat92E transcript was different from that of the protein’s, such that Stat92E protein showed clear reduction in the GB stage (Fig. S1A) and was almost non-detectable in SGs, whereas transcript decreases gradually and was still detectable in SGs, indicating that the level of Stat92E is also regulated post-transcriptionally.

Unexpectedly, other than the gradual declining of its transcription, we observed that the localization pattern of puncta of nascent transcript show a difference between GSCs and differentiating cells. In GSCs, we observed a single focus with high fluorescence intensity. In GBs and SGs, we observed two separate puncta (Fig. 1E, G). This suggested the possibility that the Stat92E locus on homologous chromosomes may be paired in stem cells, but become unpaired upon differentiation.

The observed changes of pairing pattern of the Stat92E locus prompted us to investigate whether the pairing of the Stat92E locus is under the control of early germline development and if it has any impact on the Stat92E transcript. Note that we often detected multiple (more than 2) foci of Stat92E in S-phase cells when GSC and GB are still interconnected (Fig. S1B), likely reflecting the separation of homologs or sister chromatids during DNA synthesis. Therefore, we excluded interconnected GSC-GB from our pairing assay.

### Change of pairing state on homologous *Stat92E* regions is locus-specifically regulated

To confirm the pairing state of the Stat92E locus at the DNA level, we performed OligoPaint DNA FISH [38]. Stat92E gene is located on the right arm of Chromosome3. A previous study showed that homolog pairing occurs between “button” regions in homologous chromosomes often containing a full TAD [9]. Therefore, we selected a 60Kb OligoPaint probe target region spanning the entire Stat92E gene region within a TAD (estimated based on a published Hi-C sequence data analysis (Fig. 2A, see method for determination of TAD boundary). As observed in our intron FISH experiment (Fig. 1D), the Stat92E locus showed a pattern of pairing in GSCs, detected as a single focus in a nucleus, and became unpaired, detected as two foci in a nucleus, in differentiating GBs and SGs (Fig. 2B-C, Fig. S2A-B) [39-42].

**Fig. 2.**
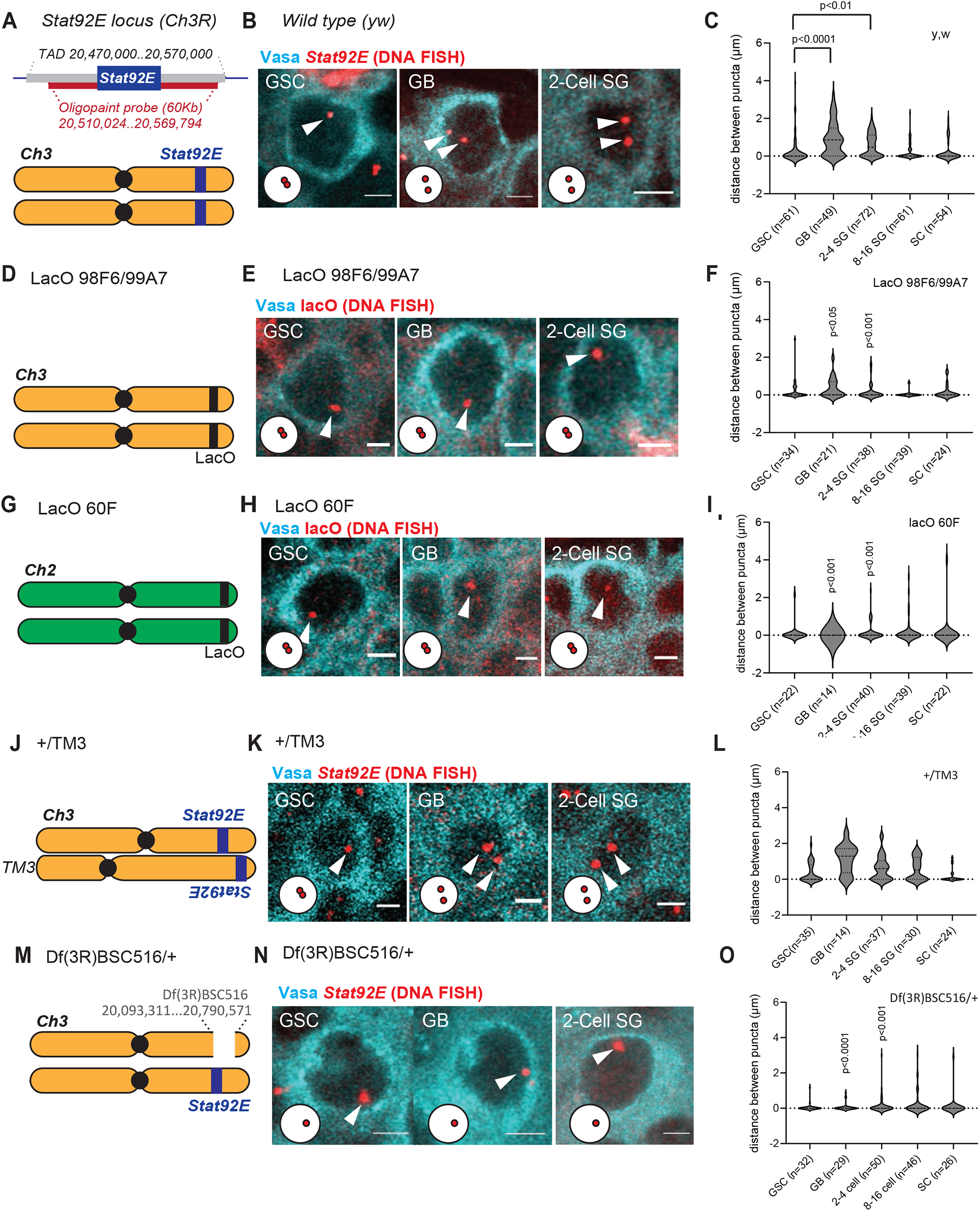
Change in pairing state of homologous *Stat92E* regions is locus-specifically regulated. **Left columns; A**) Schematic of chromosome 3 showing approximate location of the Stat92E locus. Estimated TAD boundaries and the targeted region covered by the Stat92E OligoPaint probe sets are shown at the top (see Methods for TAD boundary determination and OligoPaint probe design). **D**) Schematic of chromosome 3 showing the approximate location of the lacO 98F6/99A7 insertion. **G**) Schematic of chromosome 2 showing the approximate location of the lacO 60F insertion. **J**) Schematic showing the position of the Stat92E locus on chromosome 3 and on the balancer chromosome, TM3. **M**) Schematic of chromosome 3 showing the approximate location of the locus deleted in the Stat92E deficiency line, Df(3R)BSC516. **Middle columns; B, K, N**) Representative images of OligoPaint DNA FISH targeting the Stat92E locus in the indicated stages of germ cell development in the indicated genotypes; **B**) wildtype (yw), **K**) heterozygous for TM3 (+/TM3). **N**) heterozygous for Stat92E deficient allele (Df(3R)BSC516/+), **E, H**) Representative images of OligoPaint DNA FISH targeting lacO locus in the indicated stages of germ cells development in lacO 98F6/99A7 homozygous (**E**) or lacO 60F homozygous (**H**) genotypes. In all OligoPaint DNA FISH samples, germ cells were visualized by Vasa staining (cyan). OligoPaint DNA FISH signals are shown in red (pointed by white arrowheads). Representative pairing states are shown in lower left corner of each image. All scale bars represent 2 μm. **Violin plots (right columns);** Violin plots show the distance between puncta of OligoPaint FISH corresponding to the experiment shown in the middle panels. Cells with more than 3 puncta were omitted from the scoring (see description for Fig. S1B). Although most of cells in Df(3R)BSC516/+ flies showed only single spot plotted as distance zero (**L**), we still detected cells which had 2 puncta within a single nucleus in a low frequency, likely representing separated sister chromatids in S phase.

We next asked whether the observed pairing/unpairing events were unique to the Stat92E locus or if they occur along the entire length of homologous chromosomes. To this end, we performed DNA FISH on flies carrying an array of lacO repeats inserted at an euchromatic region on the third chromosome close to where Stat92E is located on chromosome 3 (euchromatin; Fig. 2D) or heterochromatic region near the telomere on the second chromosome (heterochromatin; Fig. 2G) and DNA FISH with oligonucleotide probes targeting the lacO repeat sequence revealed that the lacO locus on both regions remained paired throughout differentiation (Fig. 2D-I). These data suggest that the observed pairing/unpairing event seen with the Stat92E region occurs in a locus-specific manner, while other regions remain paired in these stages.

The effect of transvection depends on homolog pairing and thus is affected by chromosomal rearrangements [43, 44]. Therefore, we tested if Stat92E pairing is dependent on its position within homologous chromosomes. We still observed a similar pattern of Stat92E pairing in flies carrying the TM3 balancer chromosome (FlyBase ID: FBba0000047) in which the Stat92E locus are inverted and dislocated approximately 10Mb away from original location [45, 46] (Fig. 2J-L), suggesting that the Stat92E pairing regulation still occurs between dislocated alleles.

The observed unpairing event occurs between two homologous chromosomes and not between sister chromatids as we observed only one dot of DNA FISH signal in files heterozygous for deletion of the Stat92E locus (Df(3R)BSC516, Fig. 2M-O). This result also confirmed that the Stat92E OligoPaint probe was specifically recognizing the Stat92E locus.

The distance between Stat92E puncta progressively decreased as SGs differentiated into SCs, presumably in preparation for meiotic pairing which takes place after the SC stage [44]. Consistent with previous reports, SCs show a uniformly paired pattern in all genotypes (Fig. S2C)

### Pairing of the Stat92E locus is required for subsequent silencing of transcription

The change of locus-specific pairing state from GSC to GB prior to the downregulation of Stat92E transcription led us hypothesize that the pairing change may be required for subsequent Stat92E downregulation. To test this hypothesis, we determined the timing of Stat92E downregulation during the early stages of germline development in several genotypes. For comparison of different genotypes, we defined the ratio of mRNA levels between GSC and 2-4 SG stages, GSC/2-4SG; referred to as the “silencing index (SI)” hereafter by mRNA FISH.

Flies heterozygous for the deficiency of the Stat92E locus (Df(3R)BSC516 deleted 3R: 20,093,311..20,790,571, lacking entire TAD including Stat92E (3R: 20,470,000..20,570,000, Fig. 2A, J) must lack the effect of trans-chromosomal interaction of the Stat92E locus. Indeed, we found that there is a failure to downregulate Stat92E expression as differentiation proceeds (while in control testes the SI close to 2 (1.933), reflecting a nearly 2-fold decrease in expression as differentiation proceeds, the SI in Df(3R)BSC516/+ testes was close to 1 (1.129) (Fig. 3A, B), reflecting the unchanged expression level throughout GSC to 2-4 SG stages. Intensity of nascent transcript also showed similar level throughout GSC to 2-4 SG stages (Fig, 3C).

**Fig. 3.**
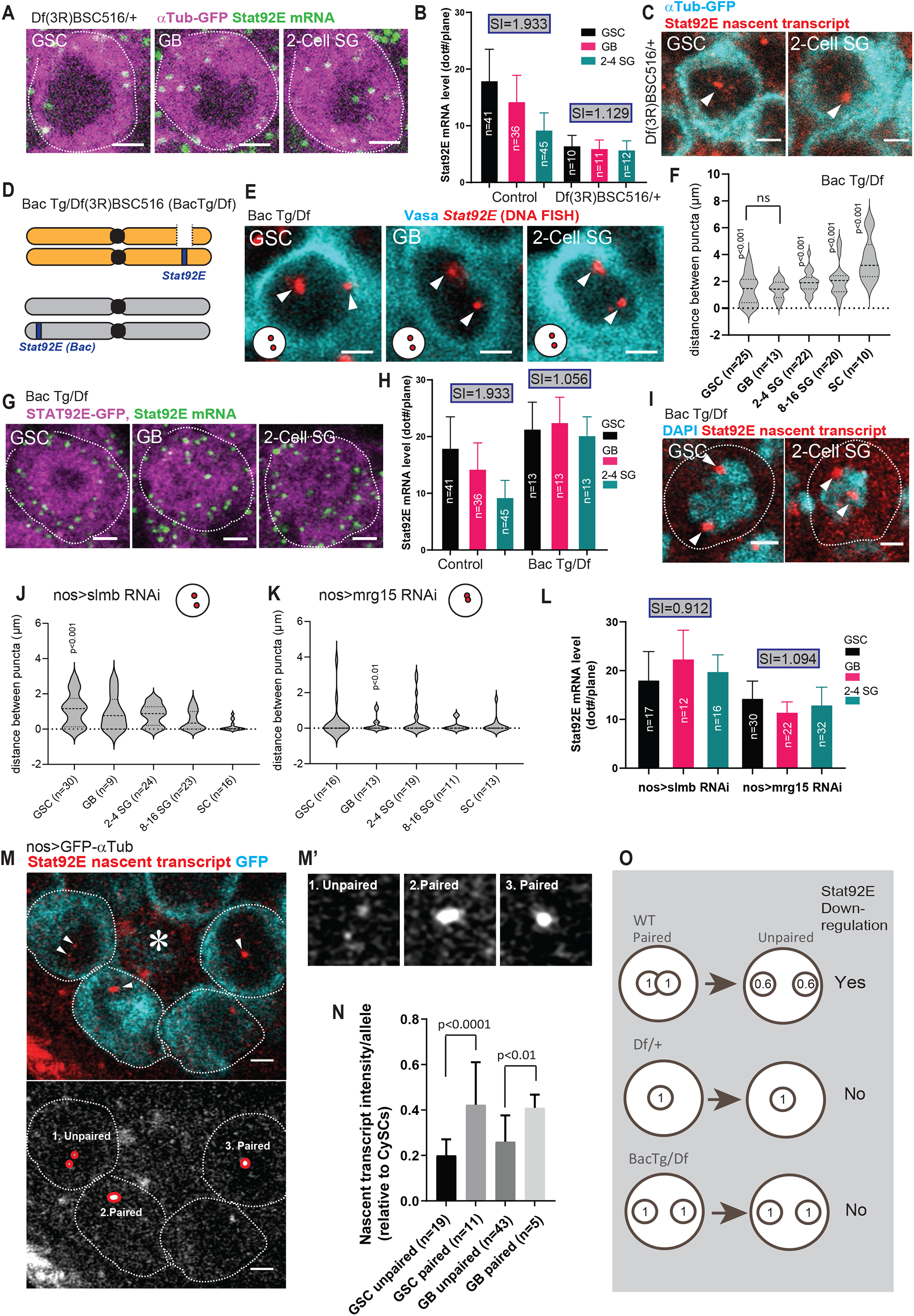
Pairing of the Stat92E locus is required for subsequent silencing of transcription. **A**) Representative images of Stat92E RNA smFISH (exon probe; green) at the indicated stages of germ cell development in Df(3R)BSC516/+ fly testes. Each dot represents a single Stat92E mRNA molecule. αTubulin-GFP (nos>αTub-GFP, magenta) was used to identify germ cells. **B**) Quantification of Stat92E mRNA level at the indicated stages of germ cell development from indicated genotypes. Y axis values are the average number of smFISH dots present in a middle plane of the cell (dot#/plane) “Silencing indices; SI” (see text) for each genotype are shown in boxes above bars. **C**) Representative images of Stat92E RNA FISH showing similar intensities of nascent transcript (intron probe; red) in GSC and 2-Cell SG in Df(3R)BSC516/+ fly testes. αTubulin-GFP (nos>αTub-GFP, cyan) was used to identify germ cells. **D**) A schematic of Bac Tg/Df(3R)BSC516 (BacTg/Df) genotype. Approximate locations of endogenous Stat92E on chromosome 3 and Stat92E Bac transgene insertion sites on chromosome 2 are shown. **E**) Representative images of OligoPaint DNA FISH targeting the Stat92E locus (red, pointed by white arrowheads) at the indicated stages of germ cell development in BacTg/Df fly testes. Germ cells were visualized by Vasa staining (cyan). Representative pairing states are shown in the lower left corner of each image. **F**) Violin plots show distances between Stat92E homologous regions at the indicated stages of germ cell development from indicated genotypes. ns: non-significant. **G)** Representative images of Stat92E RNA smFISH (exon probe; green) at the indicated stages of germ cell development from indicated genotypes. Each dot represents a single Stat92E mRNA molecule. Bac Tg contains GFP fused to Stat92E (Stat92E-GFP, magenta). **H**) Graph showing quantified Stat92E mRNA levels at the indicated stages of germ cell development from indicated genotypes (Control; nos>αTub-GFP). Y axis values are the average number of smFISH dots present in a middle plane of the cell (dot#/plane) “Silencing indices; SI” (see text) for each genotype are shown in boxes above bars. **I**) Representative images of Stat92E nascent transcript visualized by RNA FISH (intron probe, red) on Bac Tg/Df testes showing similar transcription level of both Stat92E alleles throughout all stages (GSC and 2-Cell SG stages are shown). DAPI (DNA, cyan). **J, K**) Violin plots showing distances between Stat92E homologous regions at the indicated stages of germ cell development from indicated genotypes. Representative pairing states are shown in diagrams located in right upper corner of each graph. **L**) Quantification of Stat92E mRNA levels at the indicated stages of germ cell development from indicated genotypes. Y axis values are the average number of smFISH dots appeared in a middle plane of the cell (dot#/plane) “Silencing indices; SI” (see text) for each genotype are shown in boxes above bars. **M**) Representative image of testicular niche in nos>αTub-GFP (cyan) testis. The asterisk denotes the hub area and GSCs are encircled by white dotted lines. Arrowheads indicate Stat92E nascent transcripts visualized by RNA FISH (intron probe, red), representing unpaired (1) or paired (2 and 3) Stat92E homologous regions. **M’**) Magnified images of unpaired and paired regions from (**M**). **N**) Quantification of nascent transcript intensity (measured as a ratio relative to nascent transcript intensity of CySC’s, see Method for details) in paired and unpaired conditions of GSCs and GBs. **O)** Difference in nascent Stat92E expression level between paired and unpaired conditions. Numbers in circles represent measured values of expression in **N**. All scale bars represent 2 μm.

Since Df(3R)BSC516/+ lacks one copy of Stat92E gene, other than the defective timing of downregulation, it also showed lower Stat92E mRNA expression throughout of stages (Fig. 3A-C) presumably due to having only one copy of the gene, this made it difficult to judge the downregulation timing. Therefore, we attempted to search conditions in which two copies of functional Stat92E locus are still present but they do not pair. We examined if Stat92E can pair with a Stat92E transgene located on other chromosomes and performed DNA FISH experiments of bacterial artificial chromosome transgenic (Bac Tg) line in which a 80Kb region, containing the entire Stat92E locus, is located on the second chromosome (VK00037), and introduced this chromosome into the Stat92E deficiency background via genetic crosses (Bac VK00037 Tg/Df(3R)BSC516, or Bac Tg/Df for short; Fig. 3D). In Bac Tg/Df testes, we consistently observed two spots in all cell types, representing a failure of the endogenous Stat92E gene to pair with the Bac Tg Stat92E transgene on a different chromosome (Fig. 3E-F). A previous study demonstrated that pairing occurs between separately positioned transgenes when they are contained within the same TAD [9]. The Stat92E Bac construct (VK00037) lacks ∼65Kb of the proximal portion from the predicted TAD, suggesting the requirement of this region for pairing.

Taking the advantage of Bac Tg/Df background, which does not pair but maintains the same copy number of the Stat92E gene as a wildtype, we next investigated if the regulation of Stat92E transcripts is compromised in this genotype. We examined the change of mRNA levels during differentiation in Bac Tg/Df testes and found that germ cells failed to downregulate Stat92E during differentiation, with a SI of nearly 1 (1.056, Fig. 3G-H).

There was the possibility that the Bac transgene placed in a different location may be subjected to different position effect which affected the gene downregulation timing. To exclude this possibility, we compared Stat92E nascent transcript levels between the endogenous locus and Bac Tg. We did not detect a noticeable differences of nascent transcript intensity between two puncta (Fig. 3I), indicating that the BacTg and the endogenous Stat92E locus are expressed in similar levels. Therefore, the observed defect of Stat92E downregulation in Bac Tg/Df is not due to a position effect of BacTg allele but most likely due to the absence of pairing between BacTg and the endogenous Stat92E locus.

Next, we tested the effect of global pairing and anti-pairing factors on Stat92E downregulation. Condensin II, a DNA loop extrusion factor, has been shown to antagonize homolog pairing [47-49], and its interacting factor, the *Drosophila* homolog of human MORF4-related gene on chromosome 15 (Mrg15), is known to be an anti-pairing factor [50]. The Condensin II complex is inactivated when its subunit Cap-H2 is degraded by the SCF (Skp/Cullin/F-box) E3 ubiquitin-ligase-Slimb complex. Therefore, the component Supernumerary limbs (Slmb), functions as a pairing promoting factor [47, 51]. Knockdown of Slmb in the germline significantly decreased Stat92E pairing in GSCs consistent with Slmb’s role as a pairing factor (Fig. 3J). In contrast, knock down of antipairing factor Mrg15 significantly increased Stat92E paring in GB-SGs (Fig. 3K). In both conditions, the change in pairing state between GSC and GB-SG did not occur and Stat92E failed to be downregulated during differentiation (Fig. 3L), with each genotype having a SI close to 1 (0.912 for nos>slmb RNAi, and 1.094 for nos>mrg15 RNAi).

These results so far strongly suggest that the change in pairing state promotes transcriptional downregulation. If the changes of pairing to unpairing occurs prior to Stat92E downregulation, we may detect the change of nascent transcription level between paired and unparied fraction of cells in same stages. To test this idea, we compared the level of Stat92E nascent transcript in GSCs and GBs of wild-type testes where the “change” of pairing occur most frequently (Fig. 2C), where the unpaired populations may have undergone a switch from a paired state. Strikingly, we found that the average intensity of intron signal was lower in the unpaired locus by approximately 60% of that of the paired locus (Fig. 3M-N), supporting our hypothesis that the change of pairing state of the Stat92E is important for the downregulation of its expression (Fig. 3O).

### Regulation of pairing requires Stat92E enhancer but not cis transcript

To determine the requirement of cis regulatory elements of Stat92E for pairing, we examined a Stat92E mutant allele, STAT06346 [52] in which a p-element is inserted into a putative Stat92E intronic enhancer (Fig. 4A). In the heterozygous animal of STAT06346 allele, nascent Stat92E was only detectable on a single locus, while DNA FISH shows two discrete spots in GB and SG, indicating that, as expected, the STAT06346 allele completely lacks transcription as reported previously [52] (Fig. 4B-D). In STAT06346/+ flies, we found that pairing at the Stat92E region was completely disrupted in GSCs in which the Stat92E locus is normally paired (Fig. 4E, G), suggesting that the Stat92E enhancer element is necessary for proper pairing in GSCs. The reduced expression of Stat92E mRNA in all germ cells was maintained at a similar level throughout differentiation (Fig. 4F, H) with a SI near 1 (1.172), consistent with our model that pairing is required for prompt downregulation of Stat92E.

**Fig. 4.**
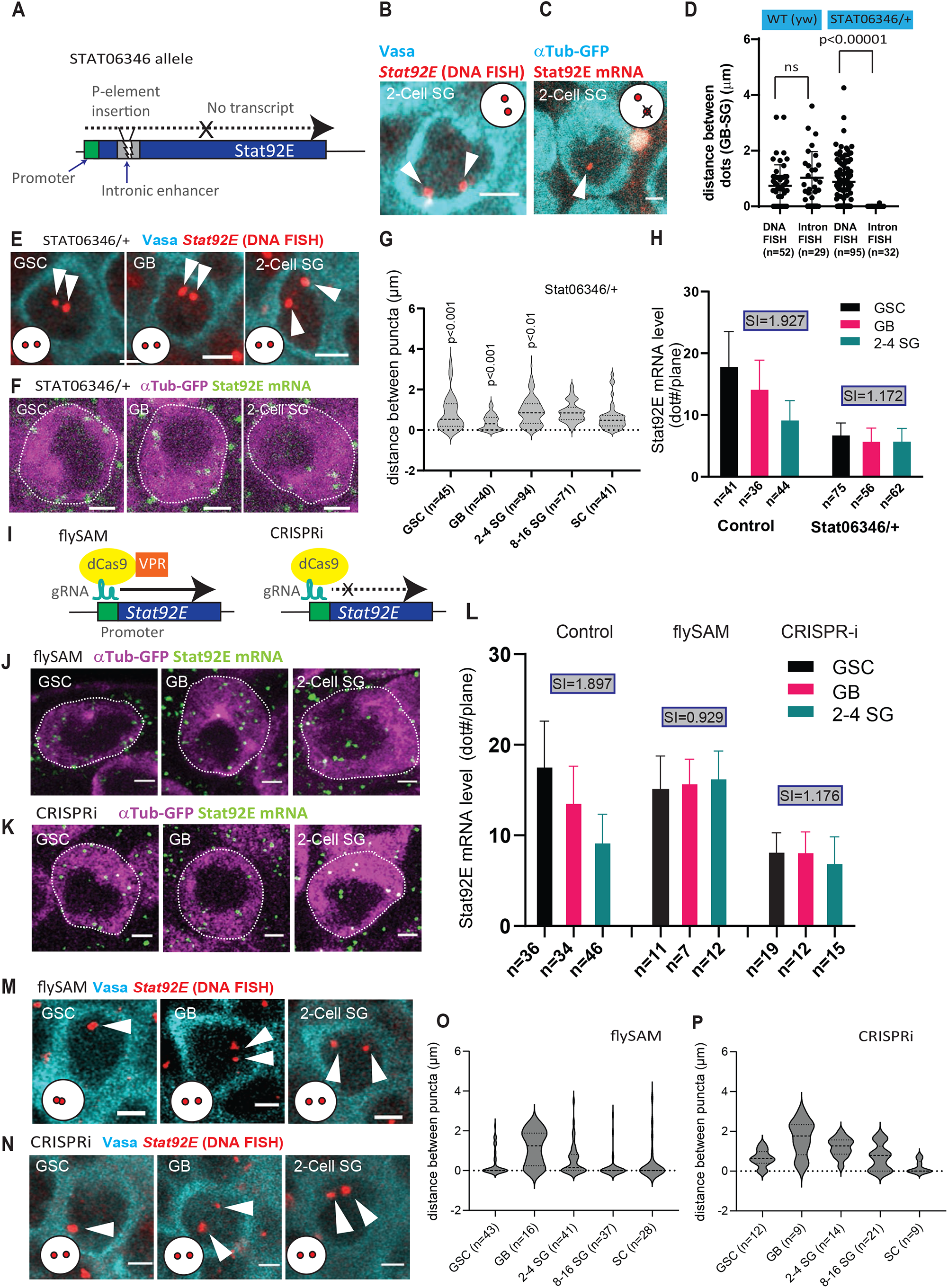
Regulation of pairing requires Stat92E enhancer but not cis transcript. **A**) Schematic of the STAT06346 allele in which P-element insertion is located in putative enhancer located in the first intron of the Stat92E gene. **B-C**) Representative images of 2-cell SGs in STAT06346/+ flies showing patterns of Stat92E DNA FISH (**B)** or nascent transcript visualized by RNA FISH with intron probe (**C**). DNA FISH showed consistently two puncta, whereas nascent transcript showed single spot, indicating that the STAT06346 allele is not expressed. **D**) Graph showing quantified distances between puncta comparing DNA FISH and intron RNA FISH between WT (y,w) and STAT06346/+ genotypes. **E**) Representative images of OligoPaint DNA FISH targeting the Stat92E locus (red, pointed by white arrowheads) at the indicated stages of germ cell development from the STAT06346/+ genotype. **F**) Representative images of Stat92E RNA smFISH (exon probe; green) at the indicated stages of germ cell development from the STAT06346/+ genotype. Each dot represents a single Stat92E mRNA molecule. αTubulin-GFP (nos>αTub-GFP, magenta) was used to identify germ cells. **G**) Violin plots show distances between Stat92E homologous regions at the indicated stages of germ cell development from the STAT06346/+ genotype. **H**) Graph showing quantified Stat92E mRNA levels at the indicated stages of germ cell development from the STAT06346/+ genotype. Y axis values are the average number of smFISH dots present in a middle plane of the cell (dot#/plane) “Silencing indices; SI” (see text) for each genotype are shown in boxes above bars. **I**) Schematics of the flySAM (left) and CRISPRi (right) systems. A gRNA targeting the transcription start site of the Stat92E gene was expressed along with dCas9. dCas9 fused to VPR artificially activates transcription irrespectively of endogenous upstream factors, whereas dCas9 without VPR (CRISPRi design) blocks endogenous transcription. **J-K**) Representative images of Stat92E RNA smFISH (exon probe; green) at the indicated stages of germ cell development from flySAM (**J**) or CRISPRi (**K**) testes induced under the nosGal4 driver. Each dot represents a single Stat92E mRNA molecule. αTubulin-GFP (nos>αTub-GFP, magenta) was used to identify germ cells. **L**) Graphs showing quantified Stat92E mRNA levels at the indicated stages of germ cell development from control (nos>αTub-GFP), nos>flySAM or nos>CRISPRi genotypes. Y axis values are the average number of smFISH dots present in a middle plane of the cell (dot#/plane) “Silencing indices; SI” (see text) for each genotype are shown in boxes above bars. **M-N**) Representative images of OligoPaint DNA FISH targeting the Stat92E locus (red, pointed by white arrowheads) at the indicated stages of germ cell development from indicated genotypes. **O-P**) Violin plots of the distances between homologous Stat92E regions at the indicated stages of germ cell development from indicated genotypes. In all OligoPaint DNA FISH samples, germ cells were visualized by Vasa staining (cyan). Representative pairing states are shown in lower left corner of each image. All scale bars represent 2 μm.

Next, we asked whether or not the effect of STAT06346 on pairing is caused by lack of its transcriptional activity. We attempted to artificially enhance cis-transcription of STAT06346 allele using the flySAM technique [53] which induces transcription by combining gRNA targeting at the Stat92E transcription start site (3R:20,552,774..20,552,796 [+]) with dCas9 fused to VPR (VP64-p65-Rta), a tripartite transcriptional activator domain (Fig. 4I). Compared to control, germline driver nosGal4-induced flySAM caused increased Stat92E expression level throughout differentiation (Fig. 4J, L). However, the pairing pattern of Stat92E was unchanged (Fig. 4M, O), suggesting that pairing regulation unlikely to be downstream of transcriptional activity.

To confirm the effect of Stat92E transcription on pairing, we next blocked cis-transcription at the Stat92E locus using the dCas9-mediated transcriptional knockdown, CRISPRi, combining dCas9 overexpression with sgRNA targeting transcriptions start site, Fig. 4I) [54]. Similar to the flySAM results, CRISPRi did not affect the pattern of pairing (Fig. 4N, P) nevertheless it caused a decrease in mRNA expression (Fig. 4K-L). These data suggest that the observed change in pairing state at the Stat92E region is not a consequence of the change in cis-transcriptional activity.

### Stat92E pairing is under the control of asymmetric histone inheritance

The observed change in pairing state was already apparent in GBs, the immediate daughter cells of GSCs, immediately the asymmetric division. During the asymmetric division of GSC, pre-existing histone H3 and H4 stay in the GSC, while newly synthesized histone H3 and H4 are inherited to the GB [21] and this pattern has been proposed to influence distinct chromatin states between GSCs and GBs. Therefore, we tested whether or not asymmetric histone inheritance contributes to the distinct pairing states of the Stat92E locus between GSC and GB. To this end, we perturbed the histone H3 asymmetry by expressing a mutant form of histone H3 that cannot be phosphorylated on its Thr3 residue (H3T3A), resulting in the random distribution of pre-existing and newly synthesized histone H3 between GSC and GB [22]. Expression of histone H3T3A-GFP in the germline under the nosGal4 driver resulted in the Stat92E locus remaining paired throughout differentiation, while the control testes expressing wild type histone H3-GFP showed the expected pairing/unpairing switch seen in wild type flies (Fig. 5A-C). nos>H3T3A did not affect the pairing states of a lacO insertion, which remained paired throughout differentiation (Fig. 5D), suggesting that asymmetric histone inheritance does not affect global pairing. The difference in number of DNA FISH puncta between nos>H3-GFP and nos>H3T3A-GFP was not due to compromised distribution of cell cycle stages in either genotype as both showed comparable frequency of 5-ethynyl-2’-deoxyuridine (EdU) incorporating S-phase cells (Fig. S5A-C).

**Fig. 5.**
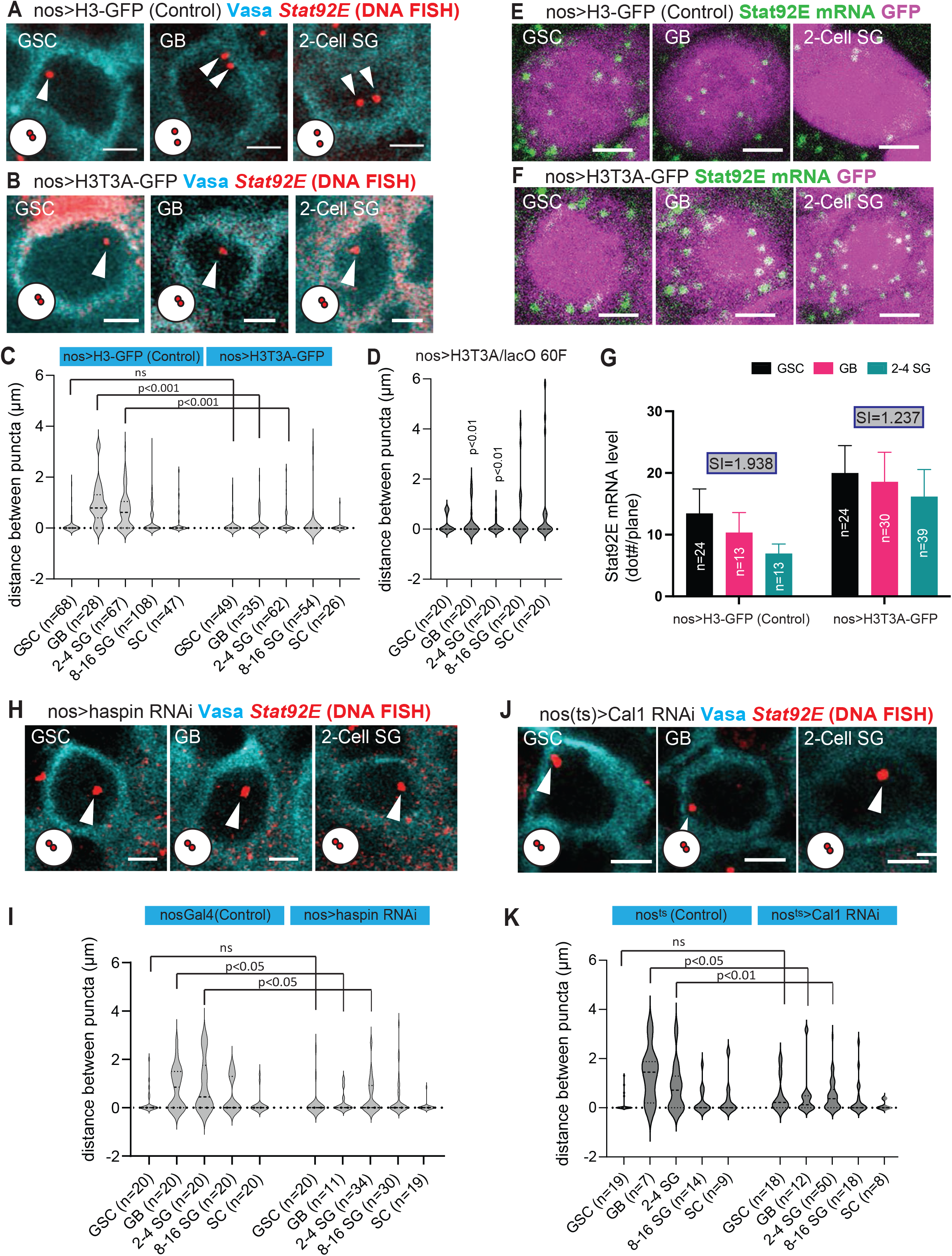
Stat92E pairing is under the control of asymmetric histone inheritance. **A-B**) Representative images of OligoPaint DNA FISH targeting the Stat92E locus (red, pointed by white arrowheads) at the indicated stages of germ cell development from Histone H3-GFP (**A**) or Histone H3T3A-GFP (**B**) expressing testes under the nosGal4 driver. **C)** Violin plots of the distances between homologous Stat92E regions at the indicated stages of germ cells from the indicated genotypes. **D)** Violin plots of the distances between the lacO 60F locus on homologous chromosomes at the indicated stages of germ cell development from lacO 60F homozygous fly expressing Histone H3T3A under the nosGal4 driver. **E-F)** Representative images of Stat92E RNA smFISH (exon probe; green) at the indicated stages of germ cell development from Histone H3-GFP (**E**) or Histone H3T3A-GFP (**F**) expressing testes under the nosGal4 driver. Each dot represents a single Stat92E mRNA molecule. **G**) Graph of the quantified Stat92E mRNA levels at the indicated stages of germ cell development from control (nos>H3-GFP) or nos>H3T3A-GFP testes. Y axis values are the average number of smFISH dots present in a middle plane of the cell (dot#/plane) “Silencing indices; SI” (see text) for each genotype are shown in boxes above bars. **H, J**) Representative images of OligoPaint DNA FISH targeting the Stat92E locus (red, pointed by white arrowheads) at the indicated stages of germ cell development from Haspin RNAi (**H**) or Cal1 RNAi (**J**) testes. **I, K)** Violin plots show distances between homologous Stat92E regions at the indicated stages of germ cell development from indicated genotypes. For Cal1 RNAi (**J** and **K**), nosGal4^ts^ driver (nosGal4 with TubGal80^ts^) was used to avoid germ cell loss. nosGal4^ts^ flies were used as control. Control and RNAi samples were both temperature shifted from 25°C to 29°C for 3 days after eclosion. In all OligoPaint DNA FISH samples, germ cells were visualized by Vasa staining (cyan). Representative pairing states are shown in diagrams located in lower left corner of each image. All scale bars represent 2 μm.

The failure to switch to an unpaired state in cells expressing histone H3T3A-GFP resulted in defective downregulation of Stat92E during differentiation (Fig. 5E-G), with a SI of near 1 (1.237), further supporting the idea that the change in pairing states of Stat92E is required for the downregulation of its expression.

To ascertain that the effect of histone H3T3A-GFP is specifically due to its effect on the asymmetric histone inheritance seen in GSC, we expressed histone H3T3A-GFP in differentiating cells using the bamGal4 driver, which is expressed later in germline development and thus should not result in perturbation of asymmetric histone H3 inheritance in GSC and GB. We performed DNA FISH on bam>H3-GFP and bam>H3T3A-GFP and found no significant change in the unpairing pattern in differentiating germ cells expressing H3T3A (Fig. S5D-F), confirming that the perturbation in pairing states seen in nos>H3T3A-GFP is due to disruption of asymmetric histone H3 distribution in GSC and GBs.

To further confirm the effect of asymmetric histone inheritance on pairing of the Stat92E locus, we knocked down two genes reported to be required for this process. Haspin is a Serine/Threonine kinase that phosphorylate Thr3 in histone H3 and its RNAi disrupts the asymmetric histone inheritance [22]. Chromosome alignment defect 1 (Cal1) is required for asymmetric sister chromatid segregation incorporated with old vs. new histone H3 and H4 in GSC and GBs [55]. Therefore, knockdown of both genes should result in the same consequence of randomized histone inheritance seen under H3T3A expression. Consistent with those results, we observed the Stat92E locus remaining paired throughout differentiation in nosGal4 driven Haspin RNAi (Fig. 5H, I) and Cal1 RNAi testes (Fig. 5J, K). Cal1 has been shown to be required for centromere pairing in meiosis [56]. However, the Stat92E locus in GB, SG stages were more paired in Cal1 RNAi, indicating that the pairing defect in Cal1 knockdown is not due to a centromere pairing defect, but likely due to a histone inheritance defect.

The niche-derived Upd signal is postulated to regulate Stat92E expression [28]. The GSC-like tumor cells induced by Upd overexpression did not show an extensive pairing pattern (approximately 50/50% of paired/unpaired pattern, Fig. S5K, L), indicating that the Upd signal is not the primary factor for dictating the Stat92E pairing change.

Taken together, our data are consistent with a model whereby the change in pairing states observed in GSC differentiation is cell-intrinsically programmed during asymmetric stem cell division, providing a new paradigm for how trans-chromosomal interaction mediate prompt gene downregulation during cell-differentiation.

## Discussion

When homologs are close together, their proximity could regulate local transcription by trans-homolog regulatory mechanisms as observed in transvection phenomena [12-17]. How the interaction of homologous regions influence local transcriptional activity and whether it occurs in endogenous gene regions have not been well understood. In this study, we demonstrate that the Stat92E gene is quickly downregulated during differentiation of the *Drosophila* male germline. The Stat92E allele is strongly paired in GSCs, and immediately becomes unpaired in GB following the asymmetric division. Disturbance of this pairing change results in a failure to quickly downregulate Stat92E expression, suggesting that the pairing change is required for switching transcriptional states. Given that enhanced (flySAM) and inhibited (CRISPRi) gene expression *in cis* did not affect pairing states, we propose that transcription is a consequence, not cause, of local pairing regulation. Finally, we show that asymmetric histone inheritance, but not niche-derived Upd signal, dictate the Stat92E pairing change, indicating that the pairing change is intrinsically programmed during the asymmetric stem cell division.

The mechanism through which Stat92E pairing change facilitates the downregulation of Stat92E expression in the *Drosophila* germline is still unknown. In conventional gene regulation, local chromatin activity is regulated by active or repressive histone marks. Histone modifiers and chromatin marks reinforce each other through various feedback mechanisms to influence local gene activity (Reviewed in [57]). One possibility is that homologous gene regions can also influence each-others’ chromatin states when they are located closely. After asymmetric division, GSC and GB are still sharing almost identical intracellular and nuclear environment. The GB is displaced away from the niche and starts receiving less signal from the niche (reviewed in [58]). The signal gradient that is initially present in the two daughter cells is quite shallow and thus how the different ‘fates’ of two daughters are ensured through differential chromatin regulation of a remains to be determined. We propose the possibility that the physical separation of homologous regions may sever inter-chromosomal interactions, which in turn may allow both alleles to be more competent to react to environmental changes. This may allow the GBs to initiate the ‘rewriting’ process of locus-specific chromatin states. Future studies will be necessary to fully understand how the interchromosomal interaction regulates the downregulation of a gene.

There is a long-standing question as to which factor stands at the top of the regulatory hierarchy and governs different cellular fates during asymmetric stem cell division. Our data suggest that programmed histone inheritance is at least one upstream factor for the change of pairing states, and the change of the pairing states may be the mechanism that can transduce the information into gene expression states. Homologous allelic pairing in a stem cell system was also reported for the *Oct4* locus in mouse, where alleles of *Oct4* transiently pair in embryonic stem cells likely to share repressive chromatin marks between homologous alleles during the transition from pluripotency to lineage commitment [57]. It is possible that pairing-dependent transcriptional regulation may be commonly used during stem cells differentiation. Comprehensive, genome-wide analysis of other gene loci will be very informative in this regard in the future.

In summary, our work provides evidence for the requirement of trans-chromosomal regulation for a switch in transcriptional state. We propose a model in which separation of homologous gene regions may be required for severing trans-homolog effect, enabling a rapid change in transcriptional activity even before intracellular (or nuclear) environment changes. Such regulation could be a conserved mechanism for prompt downregulation of gene expression status during cell differentiation.

## Acknowledgements

We thank Yukiko M. Yamashita, Michael Buszczak, Cheng-Yu Lee, Kristen Johansen, the Bloomington *Drosophila* Stock Center and the Developmental Studies Hybridoma Bank for reagents; This research is supported by 1R35GM128678-01 from the National Institute for General Medical Sciences and start-up funds from UConn Health (to M.I.)

## Author Contributions

M.I. and M.A, conceived the project, designed and executed experiments and analyzed data. M.M. and R.R. executed experiments and analyzed data. B.M. provided the methods/materials and assisted with the design of the OligoPaint DNA FISH experiments. Z.W. and S.L. analyzed published Hi-C data. All authors wrote and edited the manuscript.

## Figure legends

**Fig. S1.**
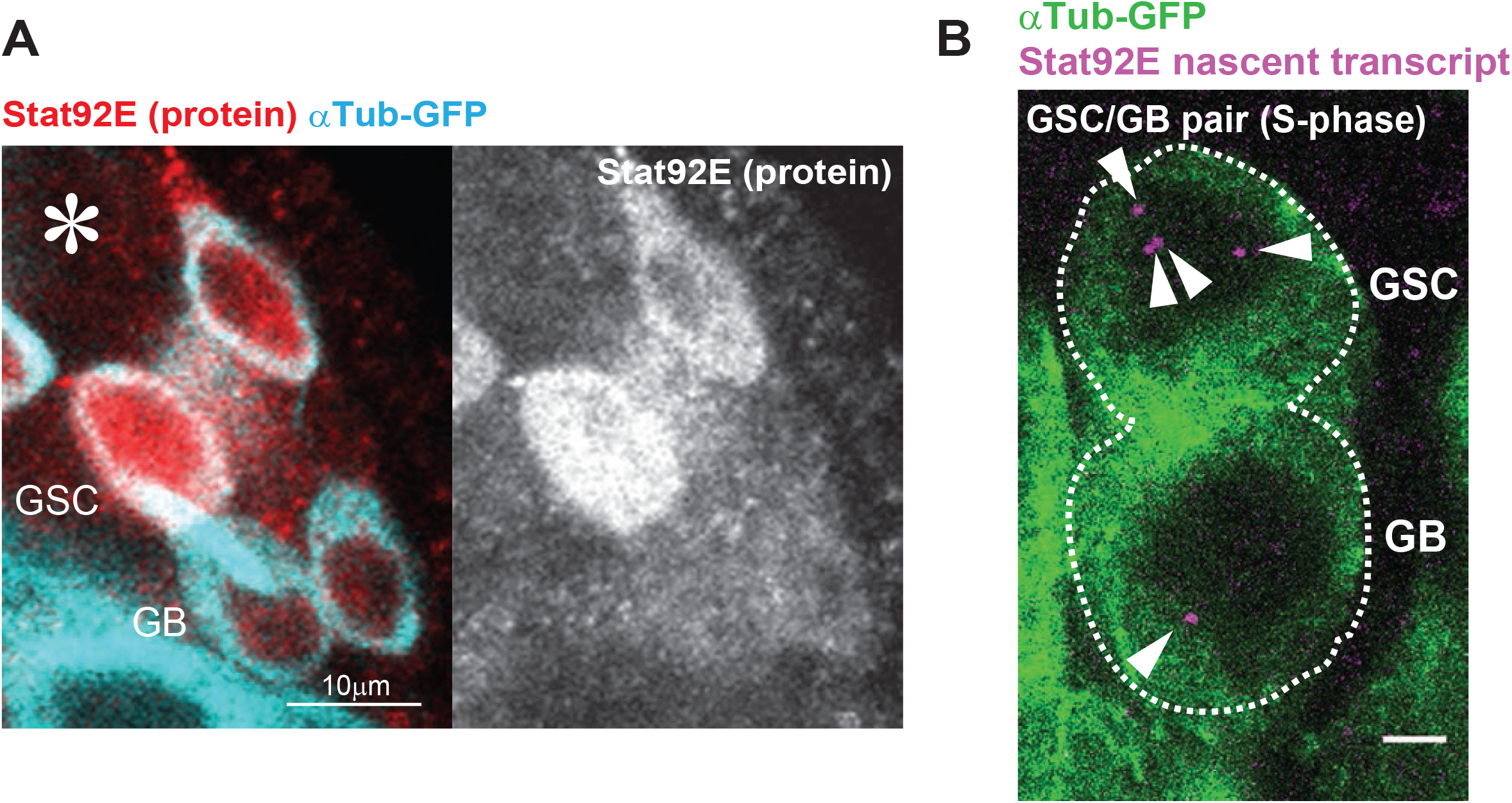
**A)** Representative image of immunofluorescence staining of Stat92E protein (red) in a GSC/GB pair in testes expressing αTubulin-GFP (cyan) under the control of nosGal4 driver. Scale bar represents 10μm. **B)** A representative image of the nascent transcript puncta (magenta, white arrowheads) in a GSC/GB pair undergoing S-phase. Germ cells are visualized by nos>αTubulin-GFP (green). Scale bar represents 2μm.

**Fig. S2.**
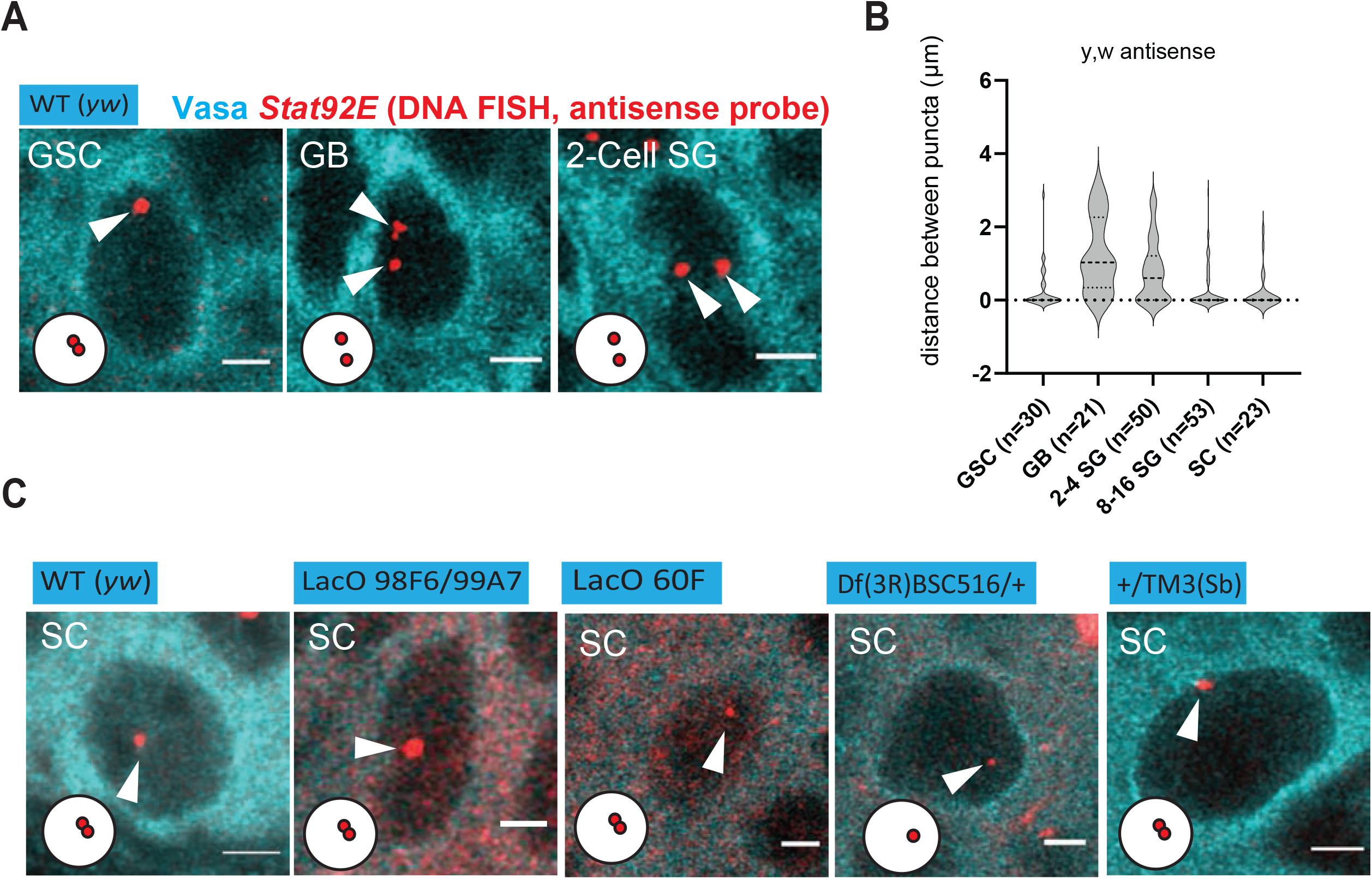
**A**) Representative images of OligoPaint DNA FISH targeting the antisense strand of Stat92E locus (red, pointed by white arrowheads) at the indicated stages of germ cell development in wild type (yw) fly testes. **B**) Violin plots showing the distance between puncta of OligoPaint DNA FISH. KDE, quantile lines, and the width of each curve correspond to the approximate frequency of data points. Cells with more than three puncta were omitted from scoring. **C**) Representative images of OligoPaint DNA FISH targeting Stat92E locus (red, pointed by white arrowheads) of SCs in the indicated genotypes. Germ cells were visualized by Vasa staining (cyan). Diagrams in the corner of images represent the pairing states of each condition. All scale bars represent 2 μm.

**Fig. S5.**
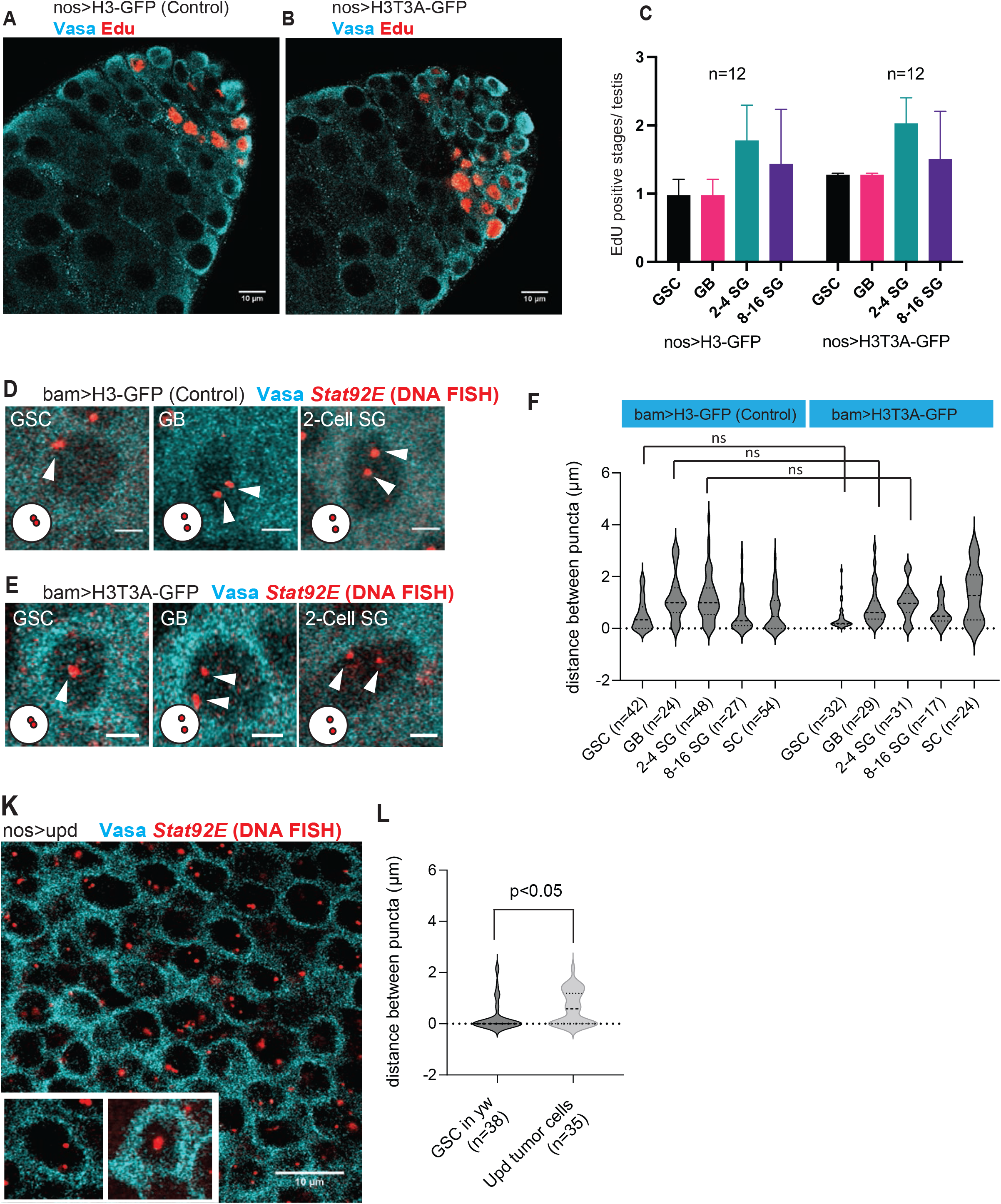
**A, B**) Representative images of EdU (red) incorporation in testes expressing histone H3-GFP (control, **A**) or histone H3T3A-GFP (**B**) under the control of the nosGal4 driver. Scale bars represent 10 μm. **C)** Graph of quantified EdU positive germ cells (or SG cysts) per testis at the indicated stages of germ cell development from histone H3-GFP (control) or histone H3T3A-GFP expressing testes under the nosGal4 driver. Because 2, 4, 8, or 16-cell SG cysts are typically synchronized to enter S-phase, we counted any Edu positive cysts as “one” for better comparison between genotypes, **E**) Representative images of OligoPaint DNA FISH targeting the Stat92E locus (red, pointed by white arrowheads) at the indicated stages of germ cell development from histone H3-GFP (**D**) or histone H3T3A (**E**) expressing testes under the bamGal4 driver. **F)** Violin plots of the distances between homologous Stat92E regions at the indicated stages of germ cell development from indicated genotypes. **K)** Representative images of OligoPaint DNA FISH targeting the Stat92E locus (red) in tumor cells of nos>Upd testes. Insets show unpaired (left) or paired (right) pattern respectively. **L)** Violin plots of the distances between homologous Stat92E regions in wildtype GSCs (yw) and nos>Upd tumor cells. All scale bars represent 2 μm unless otherwise denoted.

**Table S1**

Listed are the sequences and corresponding information for the exon and intron probes targeting an intron (Stat92E intron) and exon (Stat92E-RG) of the *Stat92E* locus. The intron probe is conjugated with Quasar 570 dye, and the exon probe is conjugated with Quasar 670.

**Table S2**

Listed are the sequences used for targeting the sense strand of the *Stat92E* locus using OligoPaint. The sequences were generated using PaintSHOP online software [58] with the dm6 genome. The oligonucleotides were flanked with common regions used for amplification by the Forward and Reverse primers and detected by the Secondary oligo (sequences shown, see *Methods* for details on OligoPaint probe production).

## Methods

### Fly husbandry and strains

All fly stocks were raised on standard Bloomington medium at 25°C (unless temperature control was required), and young flies (0-to 7-day-old adults) were used for all experiments. The following fly stocks were used: hs-flp; nos-FRTstop-FRT-gal4, UAS-GFP [59]; nosGal4dVP16 [60]; nosGal4VP16 [61]; UAS-H3-GFP [22]; and UAS-H3T3A-GFP [22]; UAS-GFP-αTubulin were gifts from Yukiko M. Yamashita; tubGal80^ts^ ([62], gift from Cheng-Yu Lee); bamGal4VP16 (gift from Michael Buszczak). lacO 98F6/99A77 (gift from Kristen Johansen).

Other stocks were from Bloomington Stock Center: lacO 60F (BDSC 25371), Df(3R)BSC516 (BDSC 25020), P[PZ]Stat92E^06346^ (BDSC 11681), Stat92E BacTg (Stat92E-GFP.FLAG, VK00037) (BDSC 38670), SAM.dCas9.GS02442 (BDSC 80517), Stat92E^TOE.GS02090^ (BDSC 80745), Mrg15 RNAi TRiP.GL00128 (BDSC 35241), Slmb RNAi TRiP.JF01504 (BDSC 31056), Haspin RNAi TRiP.GL00176 (BDSC 35276), Cal1 RNAi TRiP.HMS02281 (BDSC 41716).

### Generation of UASP-dCas9-mCherry transgenic fly

pWalium20-10XUAS-3XFLAG-dCas9-VPR vector (Addgene) was digested by NheI and SphI. dCas9 was amplified from same vector using following primers; NheI Cas9m4 F; 5’-CCATAAAACATCCCATATTCAGC-3’ Cas9m4-NLSR; 5’-AGCCCGTCCGGAACCGCTGGCCTC-3’. mCherry was amplified from pmCherry-C1 vector using following primers; mCherryF; 5’-GACGCCAGCGGTTCCGGACGGGCTGTGAGCAAGGGCGAGGAGGATAACA-3’; SphI mCherryR; 5’-GGACAGTCCTGTGCTGATATGCATGGCATGCCTTGTACAGCTCGTCCATGCCGCCGGT -3’. Obtained PCR products with digested vector were assembled by Gibson assembly (NEB) following manufacturer’s instruction. Resultant plasmid was verified by sequencing and transgenic flies were generated using strain attP2 by PhiC31 integrase-mediated transgenesis (BestGene Inc.).

### Immunofluorescence staining

Testes were dissected into 1X phosphate-buffered saline (PBS) and fixed in 4% formaldehyde in PBS for 30-60 minutes, then washed three times in PBS + 0.3% TritonX-100 (PBST) for one hour, then incubated in primary antibodies in 3% bovine serum albumin (BSA) in PBST at 4°C overnight. Samples were then washed three times in PBST for one hour (three 20 minute washes), then incubated in secondary antibodies in 3% BSA in PBST for 2-4 hours at room temperature, or at 4°C overnight. Samples were then washed three times in PBST for one hour (three 20 minute washes), then mounted using VECTASHIELD with 4,6-diamidino-2-phenylindole (DAPI) (Vector Lab, H-1200).

Primary antibodies used were: guinea pig anti Stat92E (ref, gift from Yukiko Yamashita, rat anti Vasa (DSHB).

### RNA fluorescence in situ hybridization

Fluorescence in situ hybridization was performed as described previously [63]. Briefly, testes were dissected in 1X PBS and then fixed in 4% formaldehyde/PBS for 45 minutes. Fixed testes were rinsed 2 times with 1X PBS, then resuspended in 70% EtOH, and incubated 1hour∼ overnight at 4°C. Testes were rinsed briefly in wash buffer (2X SSC and 10% deionized formamide), then incubated overnight at 37°C for 16hours in the dark with 50 nM of Quasar 570 labeled Stellaris probe against the Stat92E intron sequence and Quasar 670 labeled Stellaris probe against the second exon of Stat92E (LGC Biosearch Technologies, target sequences are provided in supplemental table S1) in the Hybridization Buffer containing 2X SSC, 10% dextran sulfate (MilliporeSigma), 1 μg/μl of yeast tRNA (MilliporeSigma), 2 mM vanadyl ribonucleoside complex (NEB), 0.02% RNase-free BSA (ThermoFisher), and 10% deionized formamide. Then, testes were washed 2 times for 30 minutes each at 37°C in the dark in the prewarmed wash buffer (2X SSC, 10% formamide) and resuspended in a drop of VECTASHIELD with DAPI.

### Detection and quantification of STAT mRNA and nascent transcript

For visualization of germ cells, all genotypes analyzed were crossed with flies expressing nos>αTubulin-GFP or nos>histone H3-GFP strains and single molecule FISH was performed using the method described above, with a probe set targeting an exon of Stat92E mRNA (Stellaris). Individual puncta represented single molecules of Stat92E mRNA [30]. Images were taken using a Zeiss LSM800 confocal microscope with a 63× oil immersion objective (NA=1.4) and processed using Zen software or Fiji. Number of puncta was manually counted per plane from an approximate mid-section of each germ cell to avoid counting the signal from adjacent CySCs.

Intron signal was measured by integrating signal from a few z-stacks and entire area of a single punctate from 0.5mm interval z-stack images in which signal was visible in 1 or 2 planes. To compare signal between samples, we normalized germline intron signal dividing by the average intron signal from 3 randomly picked CySCs located close to hub which show consistently paired, and strong signal.

### OligoPaint probe production

OligoPaint probes were designed using PaintSHOP online software [58] with the dm6 genome. The Stat92E sense probe set consisted of 949 oligos targeting the Stat92E locus and surrounding regions, from 3R:20,510,024-20,569,794 (supplemental table S2). Each oligo consisted of a region complementary to a genomic region of the sense strand of the *Stat92E* locus, flanked by a secondary recognition site (“Sec5”) on the 5’ end, and a T7 site on the 3’ end (for example: Sec5:AGCGCAGGAGGTCCACGACGTGCAAGGGTGT ttt Genomic target:ACCTGCTCCAGGTGCTTGCCGTTCTTCGGATTTatcg T7 site:tctcccTATAGTGAGTCGTATTA), (Twist Bioscience). Oligos were amplified twice by PCR (Phusion High-Fidelity DNA Polymerase, NEB) following manufactural instruction, using the following primers:

Forward: 5’-AGC GCA GGA GGT CCA CGA CGT GCA AGG GTG-3’

Reverse: 5’-TAA TAC GAC TCA CTA TAG GGA GAC GAT-3’

(Integrated DNA Technologies IDT).

PCR product was purified using Oligo Clean & Concentrator Kits (Zymo Research, D4060). RNA was synthesized from 700ng of amplified oligos using T7 RNA polymerase (HiScribe T7 kit, NEB) following manufactural instruction. RNA product was then reverse transcribed to cDNA using Maxima H Minus Reverse Transcriptase (Thermo Fisher Scientific). Briefly, 15 µl of 100 µM forward primer and 24 µl of 10mM each dNTPs (Thermo Fisher Scientific), 30 µl of 5X buffer and 57.5 µl of water were added to 20 µl of RNA product then incubated in 65 °C for 5min for denaturing. 1.5 µl of RNase out (Thermo Fisher Scientific) and 2 µl of Reverse Transcriptase were added and incubated in 50 °C for 2 hours. Template RNA was removed by alkaline hydrolysis adding 150 µl of 1:1 mixture of 0.5mM EDTA and 1M NaOH, incubated in 95 °C for 10min. Resultant single-stranded oligos were purified by using Zymo DNA concentration kit (Zymo Research, D4030) modified for short-length DNA cleaning. Briefly, 600 µl of Oligo binding buffer (Zymo Research, D4060-1-40) and 2400 µl of ethanol were added to sample then loaded onto column and followed centrifuge method in manufacture’s instruction. Purified probe was quantified with nanodrop (Thermo Fisher Scientific) and 200 pmol of probes were used for each hybridization reaction.

Stat92E antisense probes were produced using the amplified PCR product of the Stat92E sense pool as a template. Antisense oligos were amplified by PCR using oligos to add a secondary site (“Sec4”) to the 5’ end and a Sp6 site on the 3’ end. The following primers were used for amplification:

Forward: ACCCGCAGGACACCTAACCCGTCACCGTCCGATTTTTTTTTTGGAATTGTGAGCGGATAACAATT

Reverse:CCCGCAGGACACCTAACCCGTCACCGTCCGACGACTCACTATAGGGA GACGAT

The same process was followed as described for the sense pool production, using HiScribe Sp6 kit (NEB) instead of T7.

The secondary probes were designed to be complementary to the secondary sites, with fluorophores on both 5’ and 3’ ends of the oligo (IDT).

Sec4 Secondary: Cy3-TCGGACGGTGACGGGTTAGGTGCCTGCGGG -Cy3 Sec5

Secondary: Cy5-AACACCCTTGCACGTCGTGGACCTCCTGCGCTA -Cy5 Sec5

sequence: AGCGCAGGAGGTCCACGACGTGCAAGGGTGT

Sec4 sequence: CCCGCAGGACACCTAACCCGTCACCGTCCGA

lacO probe sequence: TGGAATTGTGAGCGGATAACAATT

### DNA fluorescence in situ hybridization using OligoPaint and immunofluorescence

Testes were dissected into PBS and processed for immunofluorescence as described above. After incubation in secondary antibodies, testes were washed three times (20 min each) in 0.3% PBST and then post-fixed for 10 min in 4% formaldehyde/PBS. Testes were then rinsed in 2X SSC (20XSSC was obtained from Thermo Fisher) with 0.1% Tween-20 (Thermo Fisher) (SSCT), three times for 3 minutes each. To allow a gradual transition into 50% formamide, testes were washed for ten minutes each in 20%, 40%, then 50% formamide in 2X SSCT. Testes were then heat denatured at 92°C for 30 minutes, and incubated in the probe mix at 37°C for 16 hours. The probe mix consisted of 50% formamide and 10% dextran sulfate (Sigma-Aldrich) in 2X SSCT with 200 pmol of primary oligos, 100 pmol of secondary oligos, and 1 mg/ml RNase A. The probe mix was denatured at 65°C for 5 minutes and kept on ice before adding to the samples. After the incubation, 1ml of 50% formamide/2X SSCT was added to the sample then removed with probe mix. Samples were washed again with 50% formamide/2X SSCT for 1 hour, then in 20% formamide/2X SSCT for 10min, all at 37°C. Finally, samples were washed two times, with 2X SSCT for 10min each at room temperature and mounted in a drop of VECTASHIELD with DAPI. Imaging was performed on a Zeiss LSM800 confocal microscope with a 63× oil immersion objective (NA=1.4) and processed using Airy Scan and ZEN software. Z-stacks were taken with 0.5 μm steps. Distance scoring was done using ImageJ/FIJI software.

### Detection of S-phase germ cells

S phase detection was performed using Click-iT™ EdU Cell Proliferation Kit for Imaging, Alexa Fluor™ 594 dye (Invitrogen) following the manufacturer’s instructions. Briefly, testes were dissected in PBS then transferred to Schneider’s media. 10 μM 5-ethynyl-2’-deoxyuridine (EdU) was added to the media and incubated for 2 hours at room temperature to allow EdU to incorporate. Testes were then washed three times for five minutes each in 1% BSA in PBS and fixed in 4% formaldehyde in PBS for 15 minutes at room temperature, and washed three times for five minutes each in 1% BSA in PBS. Testes were then resuspended in Click-iT™ Wash and Permeabilization Buffer for 10 minutes and then incubated with reaction cocktail for 30 minutes in room temperature. Testes were then washed three times for five minutes each in Click-iT™ Wash and Permeabilization Buffer, and fixed for 30 minutes in 4% formaldehyde in PBS then washed three times in PBS + 0.3% TritonX-100 (PBST) for one hour. Samples were then incubated in primary antibodies in 3% bovine serum albumin (BSA) in PBST at 4°C overnight and washed three times in PBST for one hour (three 20 minute washes), then incubated in secondary antibodies in 3% BSA in PBST for 2-4 hours at room temperature. Samples were then washed three times in PBST for one hour (three 20 minute washes), then mounted using VECTASHIELD with 4,6-diamidino-2-phenylindole (DAPI) (Vector Lab, H-1200).

### TAD boundary identification

We examined the published Hi-C sequencing data with topologically associating domain (TAD) coordinates in *Drosophila* larval eye discs at 10kb resolution [9] (GEO: GSE136267). TAD calls were identified by the original paper, and the details are shown below: TAD calls were based on a Hidden Markov Model (HMM) segmentation of the DI scores. The HMM was initialized with three states (downstream bias, neutral, upstream bias), each with a three-part equally Gaussian mixture model. TADs were defined as starting at the first downstream bias state following an upstream bias state with any number of intervening neutral states. We confirmed that the Stat92E gene is located within the TAD at chr3R:20470000∼20570000.

### Statistical analysis and graphing

For all violin plots, distances between two puncta of OligoPaint DNA FISH signal were plotted. Cells with more than three dots were omitted from scoring. Violin plots show KDE and quantile lines and the width of each curve corresponds with the approximate frequency of data points. P-values are comparison between each genotype with wild type data shown in Fig. 2C for each stages unless otherwise indicated. P-values are only shown for the data points which are significant (p<0.05).

No statistical methods were used to predetermine sample size. The experiments were not randomized. The investigators were not blinded to allocation during experiments and outcome assessment. Statistical analysis and graphing were performed using GraphPad Prism 9 or Microsoft excel software. Data show means and standard deviations. The P values (two-tailed Student’s t-test or adjusted P values from Dunnett’s multiple comparisons test) are provided. All experiments were repeated at least 3 times independently to confirm the result.

